# Beta2 oscillations in the hippocampal-cortical novelty detection circuit

**DOI:** 10.1101/2020.06.15.151969

**Authors:** Arthur S C França, Nils Z. Borgesius, Bryan C. Souza, Mike X Cohen

## Abstract

Novelty detection is a core feature of behavioral adaptation, and involves cascades of neuronal responses – from initial evaluation of the stimulus to the encoding of new representations – resulting in the behavioral ability to respond to unexpected inputs. In the past decade, a new important novelty detection feature, beta2 (~20 - 30 Hz) oscillations, has been described in the hippocampus. However, the interactions between beta2 and the hippocampal network are unknown, as well as the role – or even the presence – of beta2 in other areas involved with novelty detection. In this work, we used behavioral tasks that modulate novelty in combination with multisite local field potential (LFP) recordings targeting regions involved with novelty detection processing in mice – the CA1 region of the hippocampus, parietal cortex and mid-prefrontal cortex – to describe the oscillatory dynamics associated with novelty. We found that transient beta2 power increases were observed only during interaction with novel contexts and objects, but not with familiar contexts and objects. Also, robust theta-gamma phase-amplitude coupling was observed during the exploration of novel environments. Surprisingly, bursts of beta2 power had strong coupling with the phase of delta-range oscillations. Finally, the parietal and mid-frontal cortices had strong coherence with the hippocampus in both theta and beta2. These results highlight the importance of beta2 oscillations in a larger hippocampal-cortical circuit, suggesting that beta2 is a mechanism for detecting and modulating behavioral adaptation to novelty.

## 1.0 Introduction

Novelty detection is a crucial feature for behavioral adaptation and ignites cascades of neuronal responses, from the initial evaluation of the stimulus to the encoding of new representations, resulting in the behavioral ability to respond appropriately and adaptively to unexpected stimuli (Kafkas and Montaldi, 2018; van Kesteren et al., 2012). Over recent decades, an important novelty detection feature, beta2 oscillations (~20-33 Hz), has been described in the hippocampus (Berke et al., 2008; França et al., 2014; Kitanishi et al., 2015). In particular, beta2 power transiently increases during spatial novelty (Berke et al., 2008; França et al., 2014; Kitanishi et al., 2015) and its generation is implicated with AMPA and NMDA receptors plasticity between the connections of CA3 and CA1 hippocampal regions (Berke et al., 2008; Kitanishi et al., 2015). However, the interaction between beta2 with other hippocampal rhythms remains unknown. Furthermore, the hippocampus is not alone in detecting novelty: evidence in both humans and rodents points to a larger hippocampal-cortical circuit for detecting and adapting to novelty, including the mid-prefrontal cortex (mPFC) and posterior parietal cortex (PAR) (Kafkas and Montaldi, 2018; Pho et al., 2018; Spellman et al., 2015). It seems plausible that beta2 oscillations are a mechanism of communication across these regions, but there is currently no empirical evidence for or against this possibility.

Here we tested three novel hypotheses concerning the role of beta2 in novelty detection: First, whether beta2 power increase is associated with different forms of novelty (spatial and object); second, if slower hippocampal oscillations can modulate beta2 power, similarl to phase-amplitude coupling of theta-gamma oscillations during memory encoding in hippocampus,; and third, whether the novelty integration hubs in the cortex (PAR and mPFC) synchronize with hippocampal beta2 oscillations during novelty exploration.

Combining behavioral tasks where the animal is exposed to environments with different levels of novelty, and recordings from local field potential (LFP) and multi-units targeting the CA1 region of the hippocampus (HC), PAR, and mPFC, we aimed to describe the interactions among these regions involved with novelty detection processing. Using power spectral analysis, weighted phase lag index (WPLI), mean phase vector length (MPVL), Granger causality and cross-frequency phase-amplitude coupling (CFC) as indices of local and long-range synchronization (Canolty and Knight, 2010; Hyafil et al., 2015; Vinck et al., 2011), we found that transient beta2 power increases are observed only during interactions with novel contexts — environment or object — and not with familiar contexts. During novelty exploration, robust CFC was observed between theta and multiple gamma subbands. Unexpectedly, beta2 had robust coupling with delta-range oscillations. Finally, the PAR and mPFC cortices exhibited strong coherence with both theta and beta2 during novelty exploration. Within the PAR and mPFC a similar pattern of coupling between delta-ranged and beta2 was seen as in the hippocampus. The results reported in the present study also suggest that beta2 is an oscillatory feature independent of slow gamma oscillations, showing different dynamics of power and CFC, and related to novelty detection. The synchronization among hippocampus, mPFC and PAR in beta2 during novelty detection reveals its importance to understanding novelty exploration and its implication in a broad hippocampal-cortical circuit for novelty detection.

## 2.0 Methods

### 2.1 Animals

The data shown in this paper is from 9 male mice with Black57 background. All the animals were recorded in all the experimental sessions described in Figure 1A. The animals had free access to food and water. All experiments were approved by the Centrale Commissie Dierproeven (CCD) and it is according to all indications of the local Radboud University Medical Centre animal welfare body (Approval number 2016-0079).

**Figure 1.**
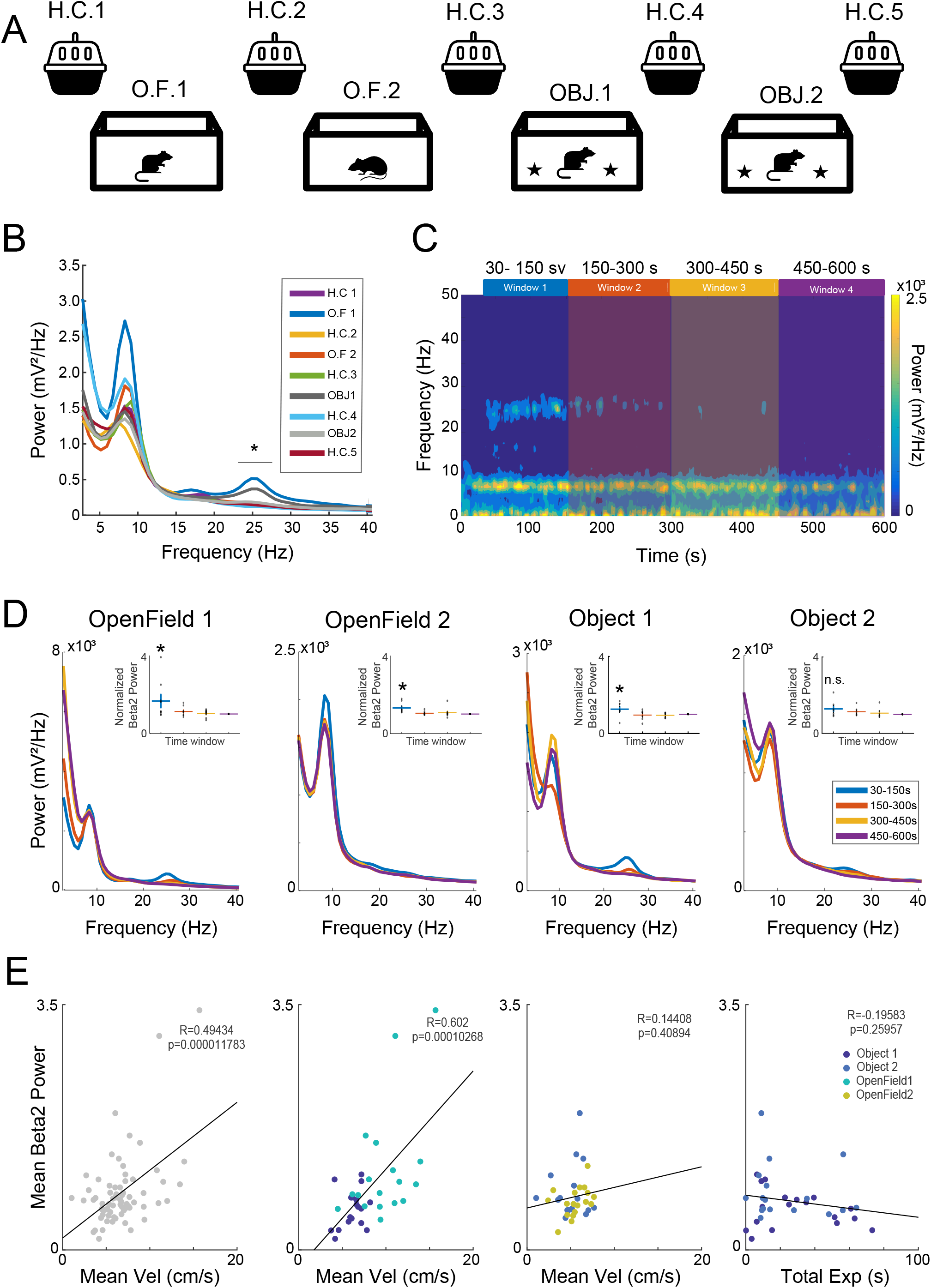
Hippocampus beta2 power increase during novelty exploration. (A) Recording sessions scheme, 10 minutes of Openfield (O.F1, O.F2) or Openfield/Object explorations (OBJ1 and OBJ2) intercalated by home cage recordings (H.C.1 – 5). (B) Group average hippocampal power spectral density over the first 150 seconds in the nine different sessions presented in panel A. Note that only OpenField1 and Object1 had increases in beta2 power, but not in other frequency bands. (C) Average of the spectrogram of the Openfield 1 session. The four time windows defined in the plot were used for all further analysis. (D) Exploration session average power spectrum density. (E) Person correlations between the mean velocity and mean beat2 power, the panel on the right correlation between total exploration time of objects and mean velocity. *p<0.05.

### 2.2 Electrode implant procedures

The self-made electrode arrays used in the present work were custom-designed to target three different regions of the mouse brain: CA1-HC, PAR and mPFC. A detailed description of the arrays and the manufacturing process can be verified at (França et al., 2020). Briefly, there were 16 channels aiming at mPFC (spread in the coordinates AP: 0.5 and 1.5; ML: 0.25 and 0.75; in three columns of electrodes in different depths - 2.0, 1.5 and 1.0), 8 channels at PAR (AP −2 and −2,25; ML: 1.0 and 1.75; DV: 0.5) and 8 channels at HC (AP −2.5 and −2,75; ML: 1.0 and 1.75; DV: 1.5).

For surgery, 10-16 week old mice were anesthetized with Isoflurane (induction at 5% Isoflurane in 0.5L/min O2; maintenance at 1-2% Isoflurane in 0.5L/min O2) [Teva]. Mice were fixed in the stereotaxic instrument [Neurostar Stereotaxic]. After shaving, the skin was disinfected with ethanol (70%). The local anesthetic Xylocaine (2%, Adrenaline 1:200.000 [AstraZeneca]) was injected subcutaneously at the incision site before exposing the skull. Peroxide (10~20% H2O2; [Sigma]) was applied to the skull with a cotton swab for cleaning and visualization of bregma and lambda. The windows in the skull through which the electrodes would be lowered into the brain were drilled specifically to accommodate the type of arrays to be implanted. To avoid contact between the dental cement and the brain, vaseline was applied to those windows after the implant. Electrodes and screws were fixated onto the skull with dental cement (Super-Bond C&B) (SuppFigure 1). Approximately 40 minutes prior to the end of the surgery, saline and analgesic (Carprofen injected subcutaneous 2.5 mg/Kg) were injected to facilitate the animal recovery.

After the experiments, animals were euthanized for post-mortem histological confirmation of electrode location. The majority of electrodes in mPFC were distributed across anterior cingulate and secondary motor cortex. The majority of the PAR electrodes were placed among layers 2 to 5. In the HC, all electrodes were located in the CA1 region, spread in different animals between stratum pyramidale and stratum lacunosum moleculare. Electrodes tracing can be verified in SuppFigure 1.

### 2.3 Behavioral task

The experiments were designed to expose the animal to different hippocampus-dependent novelty content (environment and novel object). The experiment consisted of four main different sessions of 10 min recording - two sessions at Open Field and two sessions at Open Field with Objects - interspersed by 5 minutes Home Cage recordings (see Figure 1A).

Because our goal was to evoke and investigate novelty-related oscillatory features, our task did not require or provide detailed behavioral performance output. However, to investigate how the oscillatory features investigated here were correlated with locomotor activity and behavioral exploration, the average velocity and the object exploration time were extracted. The data was computed by automated tracking of video recordings in the program Ethovision. We labeled time windows as being “object exploration” if the animal’s nose was within a quadrant draw around the object (~3cm of the object).

### 2.4 Electrophysiological analysis

#### 2.4.1 Data inspection

Electrophysiology data were acquired using Open Ephys with a sampling rate of 30 kHz. During preprocessing, data were down sampled to 1000 Hz, and EEGlab (Delorme et al., 2011) was used for visual inspection and cleaning artifacts (open channels were removed from the analysis; high frequency noises were removed by Independent Component Analysis; Segments containing large deflections in all channels were used as a criteria for recording session exclusion). Six animals had high-quality data in all recording sessions, therefore statistical analysis concerning all sessions were performed in 6 animals. Analyses in different sessions therefore have a different number of animals (varying between 7 and 9).

#### 2.4.2 Data Analysis

The data analysis was performed using custom-written and built-in routines in Matlab (R2015b). Prior to analyses, the multichannel data from each region were re-referenced to that region’s local average.

Spectral and time-frequency analysis was performed via convolution with complex Morlet wavelets (defined as a frequency-domain Gaussian with a 3 Hz full-width at half-maximum) that ranged in peak frequency from 2 Hz to 80 Hz in 100 linearly spaced steps. We reduced the dimensionality of the multichannel data by implementing a frequency-specific guided source-separation method based on generalized eigendecomposition. The goal was to create a linear weighted combination of channels (separately per region) that maximized the multivariate energy between the data covariance matrix from the narrowband filtered data, versus the broadband filtered data. This results in a single time series from each region, which was subjected to further analyses. We and others have shown that this method increases signal-to-noise characteristics while reducing computational costs and multiple-comparisons issues, and is more accurate than other source separation methods such as principal components analysis and independent components analysis (de Cheveigné and Arzounian, 2015; Cohen, 2017; Haufe et al., 2014). The Hilbert transform was then applied to these narrow-band filtered component time series in order to extract time-varying power and phase estimates.

The power spectrum was computed by averaging over the time-frequency power time series from all time points within each larger time window. Cross-frequency phase-amplitude coupling was performed in sliding windows of 5 seconds. The phase of delta-range and theta frequency (2-12Hz) and the amplitude of beta2 to mid-gamma (20 – 100 hz) was extracted. The raw CFC values were transformed into standard deviation (z) values by computing the normalized distance away from a null-hypothesis surrogate distribution, created by 500 permutations in which the phase angle time series were randomly cut and swapped. To decrease the influence of possible volume conduction we performed coherence computations utilizing weight-phase lag index (WPLI –(Vinck et al., 2011). Statistical analyses were performed using the routine RMAOV1 - Repeated Measures Single-Factor Analysis of Variance Test (alpha = 0.05).

Conditional spectral Granger causality was applied using the MultiVariate Granger Causality toolbox (Barnett and Seth, 2014). As this relies on the time-domain signals and not already-filtered data (because the causality spectrum is computed from the autoregression terms), we dimension-reduced each region using principal components analysis, taking the time series of the largest component from each region. Data were downsampled to 250 Hz and a model order ranging from 100-200 ms (varied over animals to best fit each dataset) was used for the autoregression model fitting. The advantage of the conditional Granger analysis is that it allowed us to isolate the unique contributions of one region to a second region while accounting for possible shared variance with the third region.

For detecting spiking activity, the electrophysiological signal was first band-pass filtered between 500 Hz and 6 kHz. Then, waveforms were detected using a threshold of 8 times the median absolute deviation as in Quiroga et al. (Quiroga et al, 2004) and aligned by their interpolated peak. We used the wavelet and weighted-PCA approach described in Souza et al. (Souza et al., 2019) to automatically sort the waveforms of each channel. Although we could not separate spiking activity into single units, the different MUA clusters found in the same channel presented unique activity patterns, and we therefore analyzed their activity separately. To access the phase coupling of spikes to beta2 in the first time-window (30 s to 150 s), we first selected beta2 cycles in which the mean amplitude envelope of beta2 and delta-range were both in the highest quartile. Then, for each MUA we computed the MPLV of the spikes occurring on those cycles. MUAs with fewer than 30 spikes were excluded from further analyses. The significance of each MPVL value was assessed using an equivalent surrogate distribution, computed using 500 surrogates with the same number of spikes as the original MUA. For significantly modulated MUAs we also assessed the mean phase of spiking.

## 3.0 Results

### 3.1 Beta2 power increases with both spatial and object novel content

The experiment consisted of four sessions of successive 10 min recordings. Two sessions at Open Field (OF1 Novel; OF2 Familiar) and two sessions at Open Field with Objects (OBJ1 Novel; OBJ2 Familiar) - interspersed by 5 minutes Home Cage recordings (see Figure 1A). We first verified that both spatial and object novelty evoked the same pattern of power dynamics previously described (Berke et al., 2008; França et al., 2014). As expected, beta2 power increased only during novelty (OF1 and OBJ1, Figure 1B), but not during any of the familiar contexts (Home cage, OF2 and OBJ2, Figure 1B - Table 1). We also verified the transient aspect associated with novelty-related beta2, whereby the power returned to initial levels after around three minutes of novelty exposition (Figure 1C) as previously reported (Franca et al, 2014). For all further analyses, we utilized four time windows based on the power dynamics of beta2 in the HC verified during the exploration of novelty (Figure 1C). The comparison of the normalized beta2 power (normalized by the power of the last time window) across the different time windows showed that beta2 power was higher in the first compared to the later time windows (Figure 1D - Table 1).

Next, to verify if, similar to other oscillatory activity in the hippocampus, beta2 power would be related to locomotor activity, we computed the average velocity of the animal in the different time windows defined above. Then we computed the Pearson correlation between the normalized beta2 power and animal velocity. We verified a strong correlation between the mean velocity and normalized beta2 power when looking at all experimental sessions together (OF1, OF2, OBJ1 and OBJ2 - Figure 1E). To verify if such correlation could explain by itself the previous changes in beta2 power, we computed the correlation separately in novelty (OF1 and OBJ1) and familiar sessions (OF2 and OBJ2). We found that beta2 power correlated with mean velocity in the novelty sessions, but not in the familiar session (Figure 1E), suggesting that beta2 power correlation with velocity is not general and cannot fully explain the changes seen in beta2 power. Furthermore, we verified if the amount of object exploration would be correlated with the normalized beta2 power. We found no correlation between the total time spent exploring the objects in each time window and the beta2 normalized power. At last, we found no difference (besides the window 2 and 5 of OF1) in the mean velocity between the experimental sessions (OF1 – F4,25 = 2.887, p =0.043; OF2 - F4,25 = 1.355, p =0.277; OBJ1 - F4,25 = 0.807, p =0.531 and OBJ2 = F4,25 = 1.163, p =0.352), therefore not reflecting the temporal dynamics of beta2 power. Together these results show that beta2 power increases only during novelty exposure. Beta2 has a transient aspect of higher power in the initial phase of the novelty exposure and fading towards the end of the session. The normalized beta2 power is correlated with the mean velocity of the animal exclusively during novelty exposure, but not in a familiar environment.

### 3.2 Delta modules beta2 amplitude during spatial and object novelty

Given the role of CFC in the HC during spatial navigation, learning and memory retrieval, we next explored whether there was any CFC coupling between beta2 power and the phase of slower frequencies, and if this coupling was modulated during novelty processing. We used the frequency ranges of 2-12 Hz for extracting the phase and 20-100 Hz for computing the amplitude envelope. Because the open-field exploration does not present any well-defined time event to trigger time windows for the CFC, we calculated the modulation index (MI) in sliding time windows, which allowed us to examine both the overall CFC and the temporal dynamics of CFC. We observed two key features of novelty-related CFC: First, we verified theta-gamma CFC for both low-gamma (30 - 50Hz lowG) and mid-gamma (60 - 80Hz midG); second, CFC was present between theta phase and beta2 power and between delta-range phase and beta2 and lowG power in the same time window of higher beta2 power (Figure 2A). A similar pattern of CFC was also observed in the first exploration of OBJ1 (Figure 2B).

**Figure 2.**
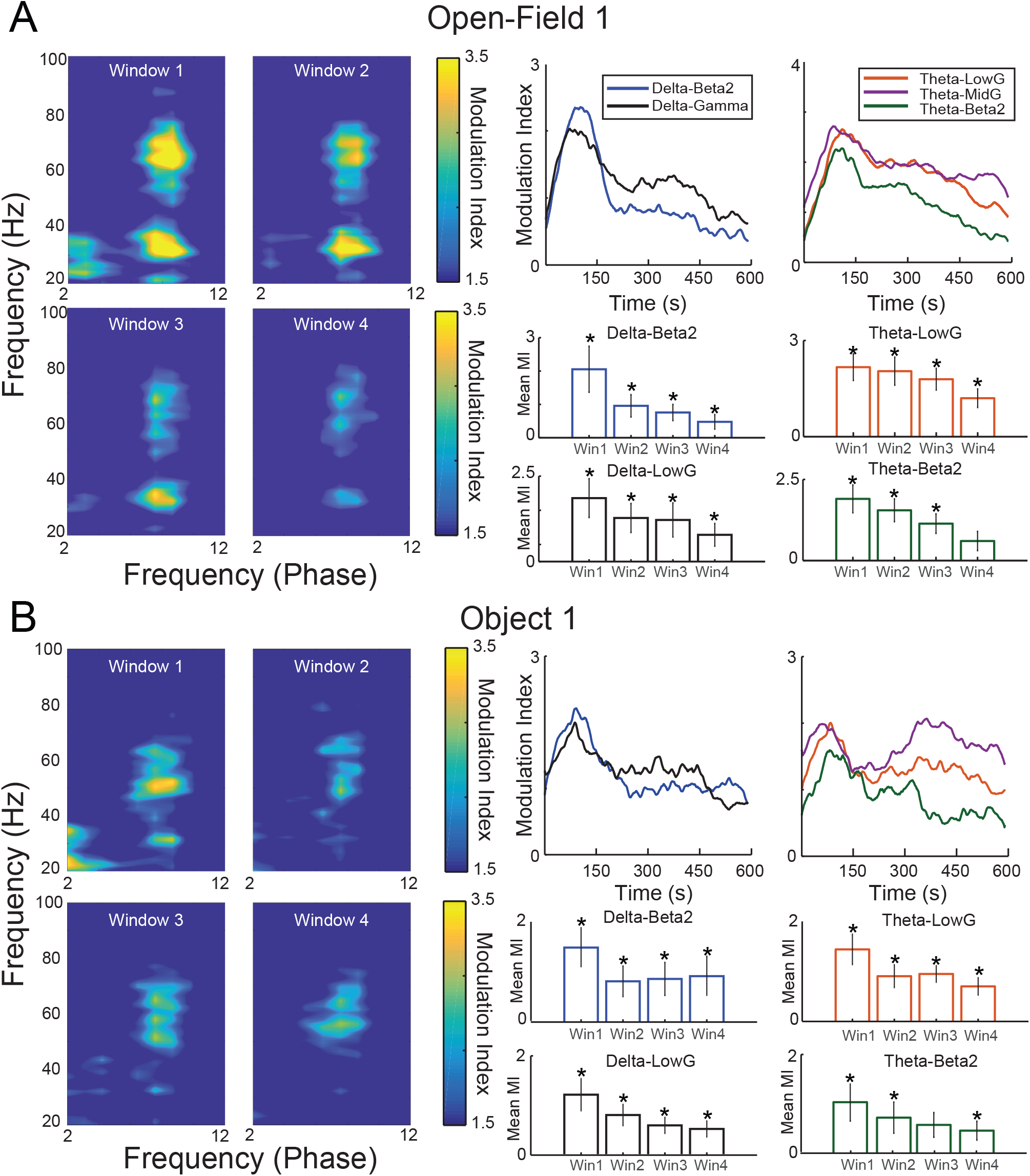
Delta-range modulates beta2 during novelty exploration. (A) Sliding-window cross-frequency phase-amplitude coupling concatenated in the four time-windows defined in Figure 1c. Right panels exhibit the modulation index during the time for different pairs of coupling. The different time-windows MI are compared against chance (0). (B) Same as (A) but for the Object 1 exploration session. *p<0.05.

Then we verified the temporal dynamics of the most prominent CFC patterns: delta-beta2, delta-lowG, theta-beta2, theta-lowG and theta-midG. For sessions with novelty (OF1 and OBJ1) the couplings of all those frequency bands were higher in the first time window, suggesting that most part of novelty detection and encoding computation happens during the initial part of the session, when beta2 power is higher (Figure 2A and B - Table 2 – time-window comparison), all pairs of coupling in all time-windows (exception to OF1 win4-theta/lowg and OBJ1 win3-theta/beta) shown MI higher than chance (Figure 2A and B - Table 2). Conversely, during the re-exposure to the Open-Field and Object (OF2 and Obj2) we observed small or no changes in the temporal dynamic of beta2 coupling (SuppFigure 2 - Table 4), but higher MI in all time-windows, exhibiting strong MI for delta-beta2/theta-beta2 and delta-LowG/theta-lowG for the OF2 session (SuppFigure 2 - Table 4). Together, these results show that besides the largely reported theta/gamma coupling, exposure to novelty is followed by delta-range/beta2 coupling with a similar transient characteristic as seen in the beta2 power dynamics, stronger MI is present in the first time window. Such temporal dynamics are not observed in the familiar sessions. However, all time-windows exhibit a MI higher than chance, indicating that beta2 events are modulated by delta-range oscillations during novel and familiar contexts.

### 3.3 Novelty modulates oscillatory coherence in hippocampal-cortical circuitry

To investigate our third key hypothesis of whether beta2 oscillations play an important role in the hippocampal-cortical novelty detection system, we computed a measure of pairwise coherence (weighted phase-lag index, WPLI) between the three regions.

We observed consistent theta-band coherence among all pairs of areas (see Figure 3A-C) in all the sessions. In addition, sessions with novelty content also exhibited increased coherence in beta2 range (Figure 3A-B). In contrast to the increase in beta2 power, this increase in coherence was not restricted to the first time window. In fact, besides the home cage 1 session (when the animal never experienced any novelty), a high beta2 coherence could also be observed even in familiar sessions, suggesting maybe the existence of a prolonged effect on synchronization among areas after the first novelty session. We also note that in the last Home exploration session the coherence between mPFC and PAR, the two cortical areas, were already in similar levels compared to the first Home exploration, while coherence between HC and mPFC, and between HC and PAR was still high (Figure 3C). This might be explained by the mechanisms underlying beta2 synchronization in the circuitry.

**Figure 3.**
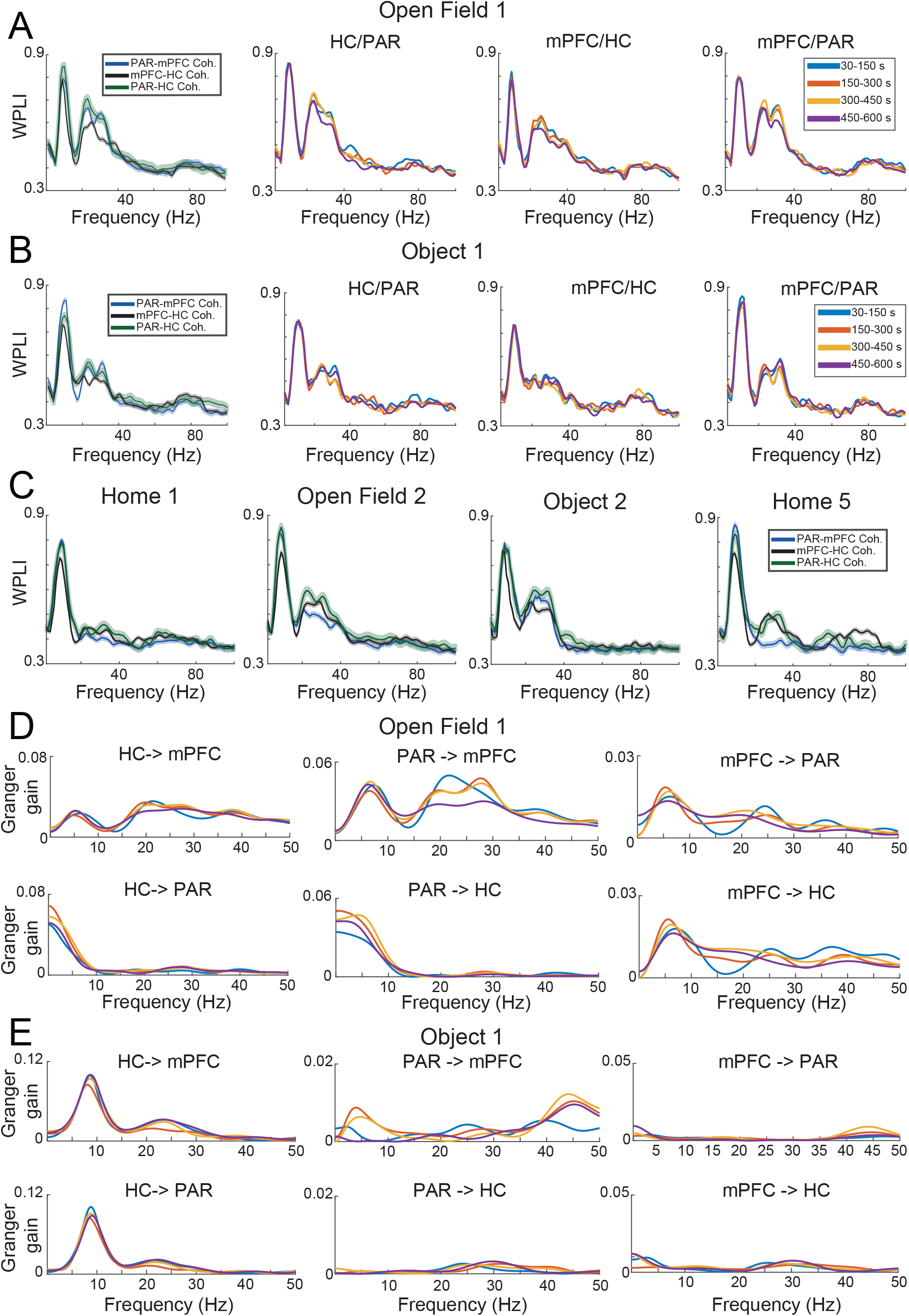
Beta2 high coherence among hippocampus, parietal and mid-frontal cortices during novelty exploration sessions. (A) Left panel shows the weight-phase lag index (WPLI) of the open-field 1 novelty exploration session. Note the high coherence in theta, beta2 and low-gamma during the first session of novelty exploration. Right panels show the different time-windows of the coherence between different pairs of regions. (B) shows the same as in (A), but for the Object 1 session. (C) Coherence plots of different familiar exploration sessions. (D) Granger causality gain between the pairs of regions in the OpenField1 session. Note the increase of granger gain in theta and beta2 range going from hippocampus and parietal cortex to mid-frontal cortex. (E) shows the same as in (D), but related to the Object 1 session.

We next applied conditional spectral Granger causality to further investigate this and to determine the causal flow of interactions around this circuit. During the OF1, PAR and HC provided input into the mPFC in theta, beta2, and lowG ranges (Figure 3D). While in the Object1 session, PAR exhibited a higher gain in the lowG frequency band (Figure 3E). HC dominated the gain values towards both cortices in both theta and beta2 (Figure 3D-E), exhibiting especially strong gain with theta and beta2 during the OBJ1 session (Figure 3E). Similar to the coherency, the Granger analysis showed no strong variation across different time windows, and suggest a less stable connectivity between mPFC and PAR cortices.

Finally, we verified the presence of beta2 burst in the raw data in all regions analyzed (Figure 4 A and B). More specifically, we could also see bursts of beta2 and lowG that happened independently (Figure 4A). To verify that cortical beta2 was not driven by volume conduction, we also analysed the multi-unit activity in the three regions of interest, computing the MPVL of the spikes in high cycles of beta2. In Figure 5A we show examples of multi-units in the mPFC, PAR and HC that are strongly coupled with the phase of beta2 (Figure 5A). We found that the multi-unit spikes couple (i.e., showed significant MPVL values) to beta2 events detected in each of the three different regions (Figure 5B-C).

**Figure 4.**
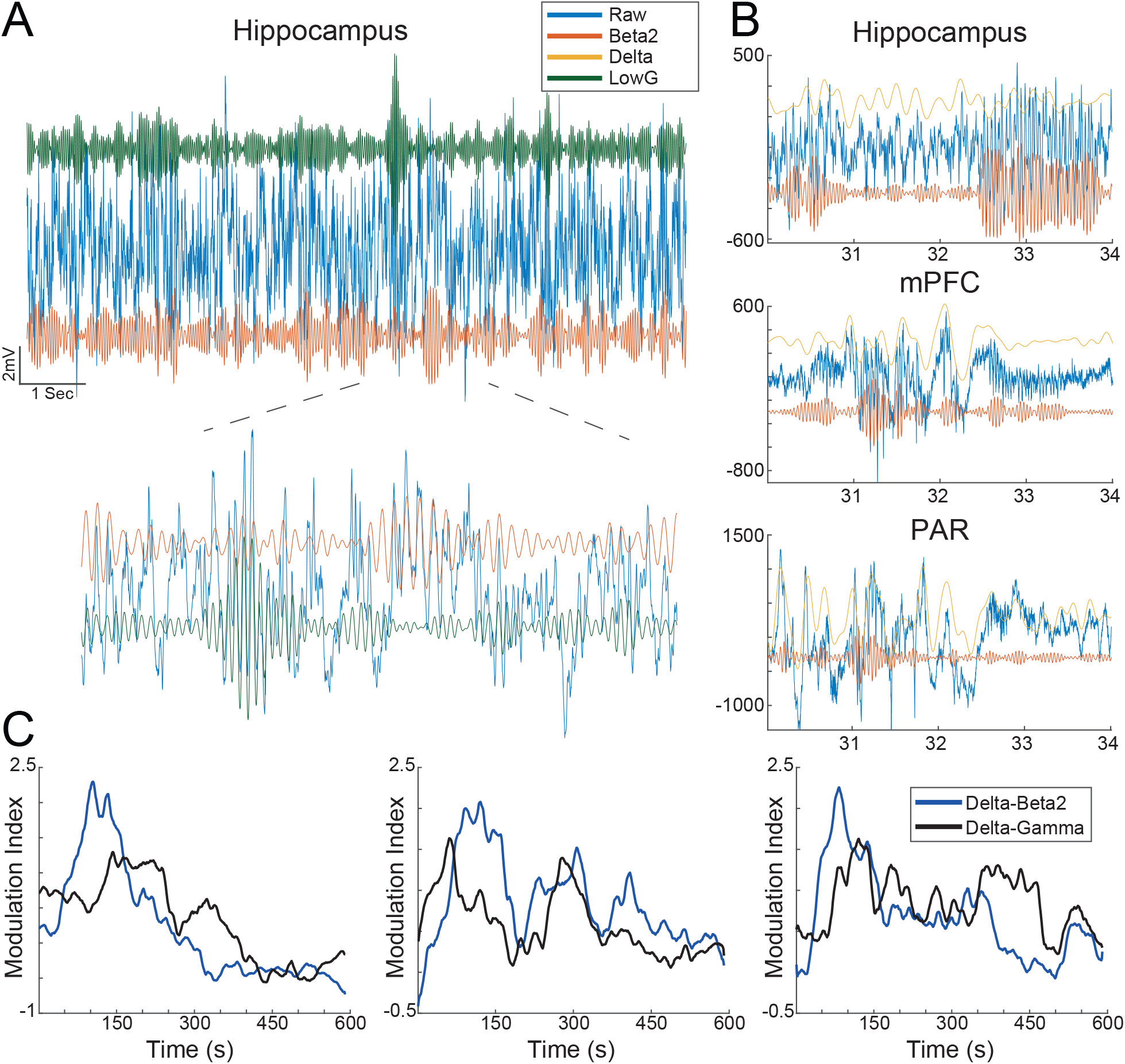
Beta2 bursts can be visualized in the raw traces of Hippocampus, Parietal and Mid-Frontal cortices. (A) Blue - Raw signal of hippocampus channel; Green – filtered signal in Low-gamma (30 – 50Hz); Red – Filtered Signal in Beta2 (20 – 30Hz). Note that the burst of Low-gamma and Beta2 happens independently from each other. (B) Blue – Raw Signal; Yellow – filtered signal in Delta (1-6Hz); Red – Filtered signal in Beta2 (20 – 30Hz). Note that The Burst of Beta2 can be verified in the Raw signal of Hippocampus, and Parietal and Mid-Frontal cortices. (C) individual examples of time-modulation index (MI) plot of Delta-Beta2 and Delta Low-Gamma during Openfield 1 exploration session. Note that the MI dynamics of Beta2 and Low-gamma are different along time.

**Figure 5.**
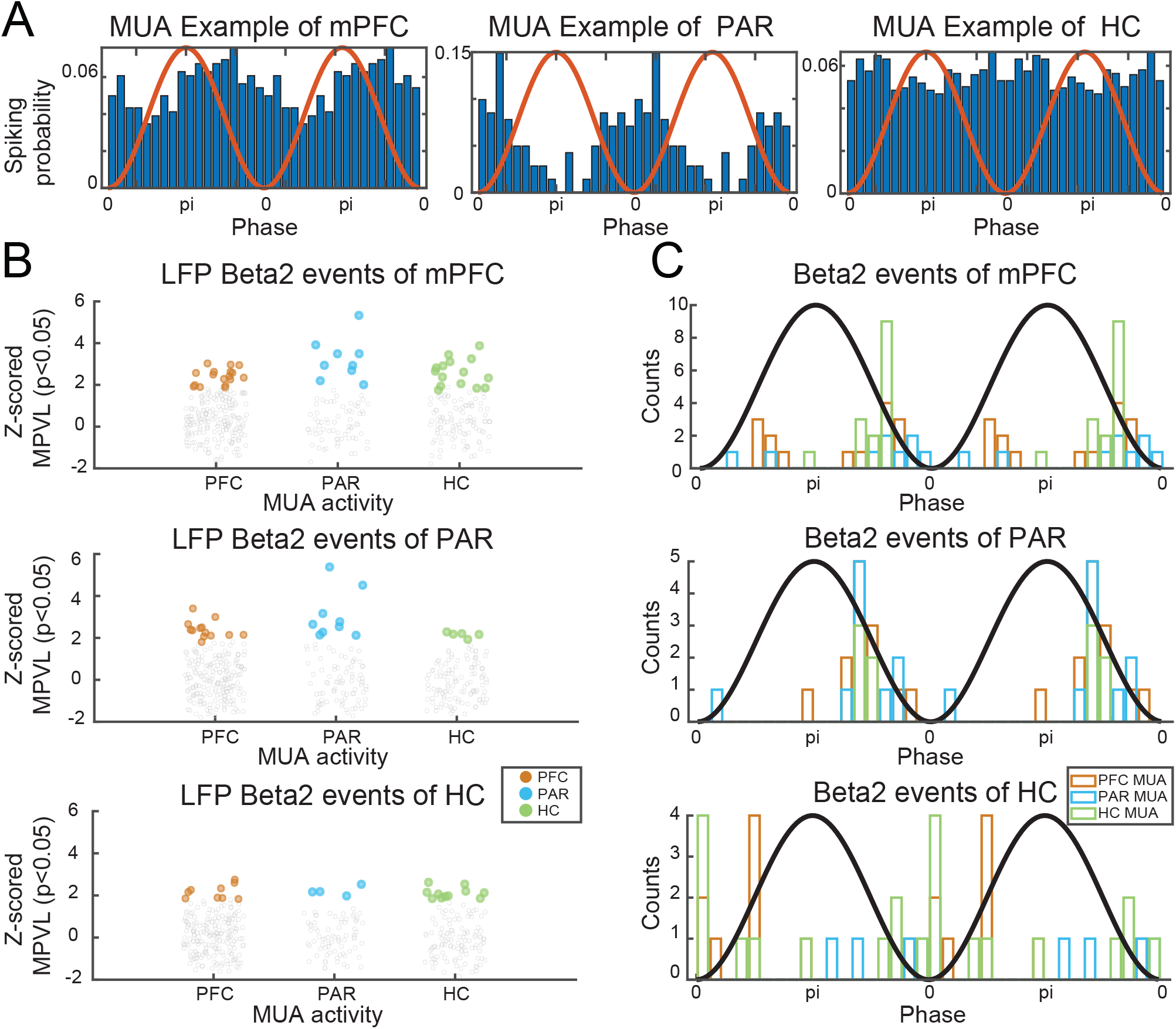
MUA coupling to beta2 events. A. Examples of MUAs in mPFC (left), PAR (middle) and HC (right) coupled to beta2 oscillations. B. Z-scored mean phase vector length for MUAs in each region in relation to beta2 events in PFC (top), PAR (middle) and HC (bottom). Colored dots denote MUAs significantly coupled to beta2. C. Histogram of the mean spiking phase of the coupled MUAs showed in B for PFC (top), PAR (middle) and HC (bottom) beta2 events. Blackline denotes the sine of the beta2.

All together, these results show that PAR, mPFC and HC synchronize in beta2, after the first novelty exposure and also in the following familiar sessions, suggesting the existence of a prolonged effect on synchronization. The contribution for such synchronization is dominated by HC towards the cortices. Finally, multi-unit activity coupled with beta2 in the three regions analyzed suggest that the beta2 events are not explained by volume conduction.

### 3.4 Parietal and mid-prefrontal cortices exhibit strong delta-beta2 coupling during novelty exploration

Lastly, to further characterize the participation of PAR and mPFC cortices in processing novelty information, we verified both power and the coupling dynamics in the cortices during novelty detection exploration. We found that the mPFC exhibited similar beta2 power dynamics as in the HC, where the increase of beta2 power was verified in the first time window. The power spectrum density (PSD) revealed an increase in beta2 frequency specifically during the novelty exploration (Figure 6A Table3). Such an increase in beta2 could be seen in the raw data, and was also independent from bursts in the HC (see Figure 4).

**Figure 6.**
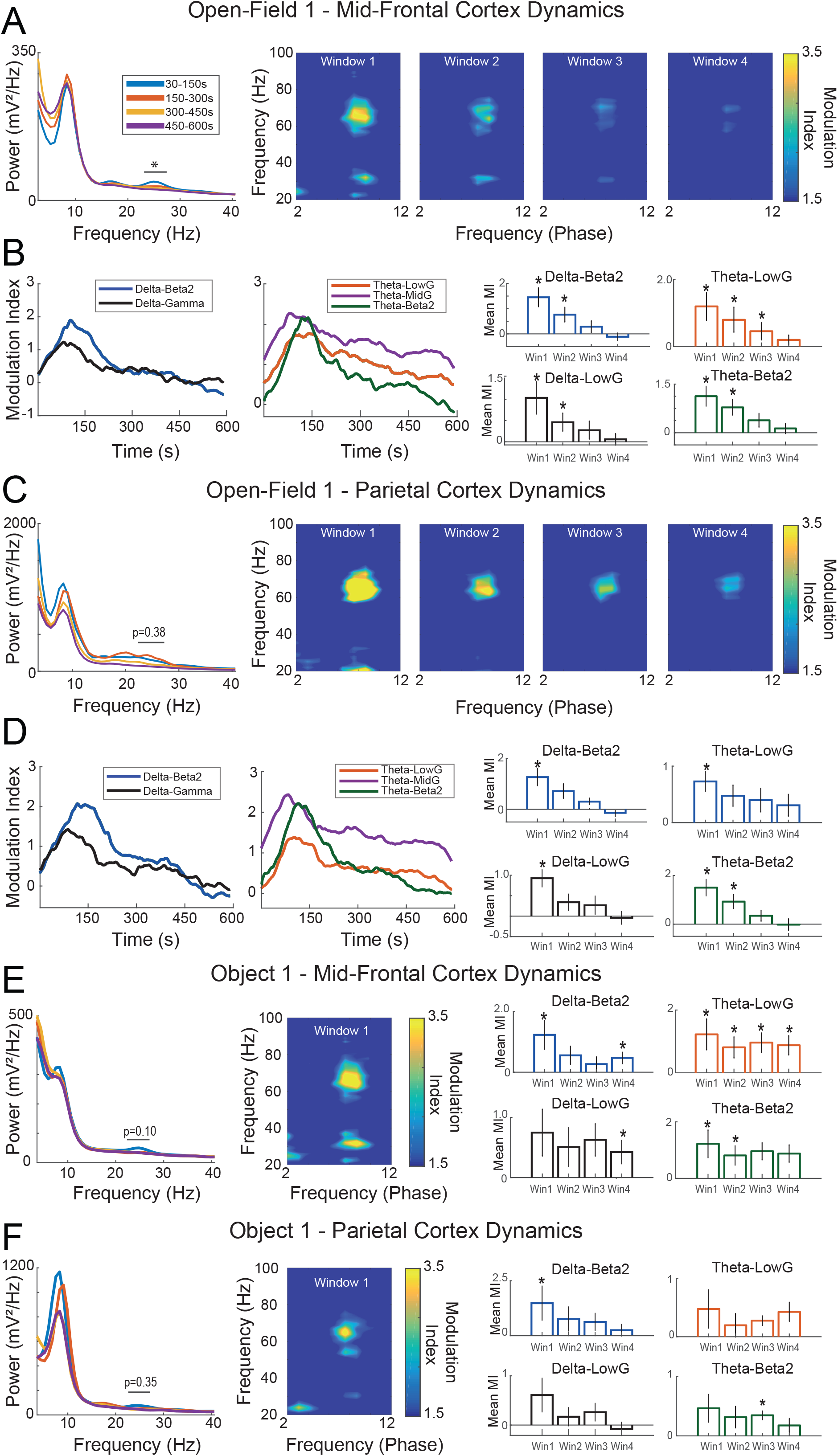
Mid-frontal and parietal cortices exhibit similar coupling dynamics as seen in the hippocampus during novelty exploration. (A) Group average Mid-Frontal power spectrum density of different time-windows. Note that the first time-window exhibits power increase in beta2 range in the first time-window. Left panels show sliding-window cross-Frequency phase-amplitude coupling concatenated in the four time-windows of mid-frontal cortex electrodes in the Open-Field 1 exploration session. Note that in the first time-window there is an increase in coupling between delta-beta2, and theta-mid-gamma, slow-gamma and beta2. (B) Modulation index during time for different pairs of coupling. The different time-windows are compared with chance (0) in the right panels. (C) Similar exhibited in (A), but for the object1 session (F 3,8 = 1.07, p = 0.38). (D) Shows the same as in (B), but for the parietal channels. Note in the first time-window the increase in coupling with beta2 and delta, and theta, but also mid-gamma and slow-gamma with theta. (D) The modulation index during time for different pairs of coupling. (E) Shows the same as in (A and B), but with Object1 session. Note the very high MI for Delta and beta2 and Mid-gamma and theta. (F) Shows the same as in (C and D), but with Object1 session. *p<0.05.

Unexpectedly, the PAR and mPFC cortices not only exhibited strong coherence among each other and the HC, but also presented a similar pattern of coupling as seen in the HC (Figure 2A-C; Figure 6 A-F Table 3), in which the novel content induced a strong modulation index (MI) between delta/beta2, theta/beta2, and theta/lowG in time-window 1. In contrast, familiar sessions did not exhibit an increase in the delta/beta2 in any time window (SuppFigure 2 B and C Table 4), in fact most of the time windows analyzed did not present any coupling during the familiar session (SuppFigure 2 and C Table 4). These results suggest that although the three areas are synchronized in beta2 during novel and familiar sessions, the coupling dynamic involving delta-range modulation is specific to novelty exposure, similar to the beta2 power dynamics. Moreover, this modulation engages both mPFC and PAR pointing to an active participation of the associative cortices in the processing information during the novelty detection sessions.

## 4.0 Discussion

In this work, we used simultaneous extracellular recordings of the HC CA1 region, PAR and mPFC to characterize, for the first time, beta2 oscillations in the hippocampal-cortical novelty detection circuit of mice. We found that beta2 hippocampal power increases during both spatial and object novelty, but not during the exploration of familiar contexts. We have shown that delta-range oscillations modulate beta2 and lowG during the exploration of new and familiar environments, while theta modulates beta2 and lowG and midG. In addition, we found strong coherence in theta and beta2 bands during novelty exploration among the areas recorded, in which the higher Granger gain for beta2 and theta came mostly from HC. Such coherence was translated into the increase of beta2 power in the mPFC but not in the PAR, even though bursts of beta2 could be identified in the raw trace of both mPFC and PAR, as well as beta2-modulated multi-units. Finally, we have observed similar coupling characteristics in the cortex to what is described in the HC, showing that beta2 is also modulated by delta-range activity in the cortex. Taken together, these results highlight the importance of beta2 oscillations in a larger hippocampal-cortical circuit, suggesting that beta2 is a mechanism for detecting and modulating behavioral adaptation to novelty.

The three regions investigated in the present study, HC, PAR and mPFC, 1 - have monosynaptic connections among each other (Cenquizca and Swanson, 2007), 2 - are extensively related to learning and encoding of memory (Hasselmo, 2006; de Lima et al., 2011; Lisman and Grace, 2005; Lisman and Otmakhova, 2001, Preston and Eichenbaum, 2013; Sigurdsson et al., 2010; Spellman et al., 2015, Cross et al., 2013), a characteristic which is preceded by novelty detection (van Kesteren et al., 2012) and 3 - are implicated in novelty detection networks in human models (Kafka and Motaldi, 2018).

To coordinate the activity of such diverse brain areas during the process of novelty detection, oscillations are suggested to play a key role in the integration and coordination of the information (Buzsáki and Draguhn, 2004; Fries, 2005). Theta oscillations are thought to coordinate neural networks during memory encoding within and across different areas (Benchenane et al., 2011; Colgin, 2015; Tort et al., 2009). The close relationship between HC and mid-prefrontal areas as it relates to memory encoding and retrieval has been extensively reported (Benchenane et al., 2010, 2011; I.b.h, 2019), and theta plays an important role in mediating the function of these two areas (Benchenane et al., 2010, 2011). But until now, no specific oscillatory dynamic responsive to novelty content was reported playing a role in the coordination of different brain areas responsible to process the novelty information.

We and others have identified beta2 as an oscillatory feature in novelty detection in the HC (Berke et al., 2008; França et al., 2014; Kitanishi et al., 2015). This oscillation has a spectral peak around 20 to 30Hz in mice (Berke et al., 2008; França et al., 2014), and slightly faster in rats – 25 to 48Hz (Kitanishi et al., 2015). The beta2 is elicited with spatial/environmental novelty (Berke et al., 2008; França et al., 2014; Kitanishi et al., 2015), but not with olfactory novel stimulus (Berke et al., 2008). Beta2 also has transient characteristics, it reaches the peak during the first 2 minutes after the novelty presentation and decreases with time (Berke et al., 2008; França et al., 2014). It is suggested to originate in the projections of CA3 towards CA1 (Berke et al., 2008; Kitanishi et al., 2015). Beta2 has been related to the stability of place fields (Berke et al., 2008; Kitanishi et al., 2015), memory consolidation impairment of novel recognition (França et al., 2014) and implicated to drive the synaptic delivery of GluR1-containing AMPA receptors (Kitanishi et al., 2015) and CA3 NMDA receptors (Berke et al., 2008;). Finally, beta2 has been reported absent in primary sensory and motor cortices (França et al., 2014), regions not associated with novelty detection (França et al., 2014).

In the present work, we replicate the main features in the power dynamics reported before (Figure 1A-C, Table 1). Similarly to the previous reports, we have shown that beta2 can be verified at the raw signal of the HC channels (Figure 4). The results reported in the present work, in accordance with what has been previously described, shows a delay between the beginning of the novelty exposition and the peak of beta2 (Figure 1C, Table 1). This latency period may reflect the generation of a mismatch from previous expectations (Grossberg, 2009) or the time that animals take to perceive the experience as novel. Another possibility is the delay being related to the stability of the place field that is followed by the dynamic of beta2 (Berke et al, 2008). We also replicated the relation between beta2 normalized power and the mean velocity of the animal (Figure 1E - França et al 2014); this correlation is only present in the novelty exposition sessions, and not in the familiar exploration sessions (Figure 1E). As previously reported, no correlation between object exploration and beta2 normalized power was found (Figure 1E - França et al 2014). Because mice have an innate exploratory behavior when they are exposed to novel environments, it is expected to see an increase of the total distance traveled and thus the mean velocity in novel environments. However, except for one pair of time-windows in OF1 (2nd and 5th windows, in which the animal should be more habituated to the novelty), the mean velocity did not statistically change within the exploration session, while beta2 power varied along the session (Figure 1) suggesting that the correlation with velocity might reflect the behavior output expected of novelty sessions, as opposed to velocity directly driving beta2 activity.

One of the novel results reported here was the cross-frequency modulation between a slow frequency range within the delta-range activity and the power of beta2 during novelty detection. This set of results was surprising, and not anticipated for the experimental design. As recently shown, delta oscillations have been related to the respiration rhythm (Lockmann and Tort, 2018; Tort et al., 2018). However, the only way for checking if the phase of the slow oscillation reported here is indeed related to a delta oscillation was implanting electrodes in the olfactory bulb. Therefore, the results present here were reported as a delta-range oscillation and future research is needed to further investigate the relation between respiration and novelty detection.

The sliding time-window CFC analysis in the HC, especially during the first 150 seconds (window 1), revealed theta-nested spectral components, consistent with previous reports (Lopes-dos-Santos et al., 2018). The CFC revealed the peak of theta/beta2 around 22Hz (instead of the 25Hz of beta2 power increase – Figure 2A), Theta/lowG at 35Hz and theta/midG around 70Hz (Figure 2A) or 54Hz (Figure 2A). We observed an increase of theta/midG coupling [1] during the exploration of novel environment and objects (Figure 2A and B), while theta/lowG coupling was more prevalent in the “retrieval” at the familiar session (SuppFigure 2A Table4) in accordance with previously reported theta/midG coupling increases during learning and retrieval of memory (Lisman and Jensen, 2013; Lopes-dos-Santos et al., 2018; Tort et al., 2008, 2009; Zheng et al., 2016). Interestingly, beta2 and lowG were strongly modulated by the delta-range phase (Figure 2A and B). Although this could initially point to beta2 and lowG as being part of the same oscillatory regimen, beta2 and lowG have different spectral peaks (25Hz vs 35Hz), beta2 has a transient power characteristic and lowG does not (Figure 1 C, table 1) and inspecting the raw signal reveals that these two dynamics can be observed independently of each other (Figure 4A). Furthermore, beta2 and lowG have different coherence peaks (Figure 4) and exhibit different temporal coupling dynamics during novelty exploration (Figure 4C). The usage of the same nomenclature (lowG to describe beta2 and lowG) may create difficulties in the characterization of the function behind these different oscillations, which could also be the reason for beta2 being reported only twice in the past decades (França et al., 2014). Instead of only the band of frequency, in which authors constantly change the frequency range for the same nomenclature, the oscillations ideally should be classified based on different characteristics, from the species been recorded to wave-shape, origin and physiological function (Cole and Voytek, 2017; Tort et al., 2018).

The distinction between beta2 and lowG is also important in the perspective of a complex network involving different brain regions because beta oscillations are implicated in long-range synchrony between different areas of the brain, a feature not shared with gamma oscillations (Kopell et al., 2000). For the first time, we revealed that during the novelty exploration sessions the HC has a strong coherence in the beta2 frequency band (Figure 3A and B), such coherence is not seen when the animal never faced the novelty content before (Figure 3C). On top of that, Granger causality revealed that the highest Granger gains come from HC and PAR cortex towards mPFC in theta, beta2 and lowG frequency band during OF1 exploration, while in the object novelty session the Granger gain comes mostly from HC (Figure 3D and E). Note that during the subsequent familiar exploration sessions the beta2 coherence in all three areas remains strong, probably carrying novelty content information towards the cortices, which may act as hubs for comparing the familiarity/novelty contents. In contrast to the beta2 power dynamics, which increase only at the beginning of novelty sessions, this suggests a more cumulative effect on coherence. We also notice that in the last HC session, the coherence between the two cortices decreased while their coherence with the hippocampus was still high. This might be explained by the strong hippocampal influence in the generation of beta2, or memory trace retrieval characteristics previously described between mPFC and HC (Jin and Maren, 2015). Further investigation is needed to reveal detailed aspects of these interactions. In summary, the coherence and Granger results presented here point to the close communication among the three areas recorded, showing that all three areas communicate via theta and beta2 during novelty and familiarity exploration.

We also have shown that beta2 has similar transient power dynamics also in the mPFC, increasing at the beginning of the session and fading towards the end of the session (Figure 6A). Although PAR did not exhibit a statistically significant increase in beta2 power, the beta2 bursts can be verified in the raw signal of mPFC, and PAR LFP was coherent with other areas (Figure 4). Furthermore, all three areas involved showed multi-unit coupling with beta2 bursts, including PAR multi-unit activity coupled to the beta2 bursts of mPFC and PAR (Figure 5). This corroborates the results of coherence and Granger causality analyses, showing that the cortical beta2 is not a result of volume conduction from the hippocampus. Lastly, we have shown that similar to HC coupling dynamics, 1-both cortices exhibit strong coupling between theta/midG and theta/lowG during novelty exploration; and 2- that both cortices show the same coupling between delta/beta2 as exhibited in HC in the first time window that beta2 exhibited higher power (Figure 6B, D, E and F). These couplings are only found during the novelty exploration (Figure 6), and not during familiar exploration (SuppFigure 2 B and C). Even though there is a trend in the delta-beta2 coupling to be higher in the first time-window, this effect is stronger in the OF1 session (time-window effect in mPFC, PAR and HC; see Table 2 and 3). Thus it’s not clear whether this modulation follows the temporal dynamics of beta2 power, coherence, or a mix between them. Similar coupling was previously reported in the mid-prefrontal cortex during recording in freely behaving rodents (Andino-Pavlovsky et al., 2017) or during learning and working memory (Canolty and Knight, 2010; I.b.h, 2019). However, for the first time we show that the local delta oscillations modulate the beta2, not only in the HC, but also in the PAR and mPFC during novelty exploration.

Importantly, beta2 coherence among the three areas showed a temporal dynamic different from beta2 power, with a long-lasting effect across even sessions. This unveils the existence of multiple processes influenced by beta2 oscillations: one in a shorter timescale, revealed by the transient presence of hippocampal beta2 bursts during novelty exposure; and another, in a longer timescale, is characterized by the beta2 synchrony across the hippocampus, mPFC and PAR that in our data extends through the entire session of novelty exposure and even further into familiar sessions. Those two mechanisms might be associated with different steps of memory encoding. For example, Grossberg (2009) suggests that initial beta2 bursts could be a mechanism for the fast stabilization of the memory traces (during memory acquisition), explaining the rapid emergence of place cells in the HC (Grossberg, 2009; Berke et al., 2008). It has also been shown that inhibition of protein synthesis in the HC impairs reconsolidation of memory traces only when the memory reactivation involves novelty (Rossato et al., 2007; Radiske, A. et al. 2017) - that is, in the presence of beta2 bursts. Both of those processes, memory acquisition and reconsolidation, involve first setting the memory into an active state, which requires further stabilization towards an inactive memory state (Nader 2015). Thus, there might be a link between the acquisition/activation of memory traces and the initial beta2 bursts. On the other hand, beta2 coherence between HC and the two cortices stays higher for a longer time after novelty exposure, which could indicate a role in the stabilization of the memory traces and the LTP induction that happens in the HC (Clarke, J. R. et al., 2010). Despite this being an interesting hypothesis, new experiments are needed to specifically investigate the direct relation of beta2 to the different memory trace processes.

Finally, in between these two temporal dynamics of beta2 there is the modulation of beta2 amplitude by delta-range oscillations, which seems to follow a short timescale in the cortex only during novelty, similar to the transient beta2 bursts, and a longer timescale in the HC, even though the modulation tends to be higher in the first time-window.

Together, these results highlight and further support the relation of beta2 oscillations and novelty - extending it to a larger hippocampal-cortical circuit - and suggesting beta2 as a mechanism for detecting and communicating information among the areas involved in behavioral adaptation to novelty.

## Acknowledgments

MXC, NB and ACSF are funded by ERC-StG 638589. BCS is funded by NWO TOP grant 612.001.853.

---

**Supplementary Figure 1 – Electrodes positioning.** (A) shows an example of electrode implant surgery. Left panel the two craniotomies for the electrodes of parietal cortex and hippocampus, and for the mid-frontal electrodes. Right panel shows the cap holding the implant to the head of the animal. (B) Examples of electrodes tracks found in mid-frontal cortex of the animals. (C) Example of electrodes tracks found in the parietal cortex. (D) Example of electrodes tracks found in the hippocampus.

**Supplementary Figure 2 – Cross-Frequency coupling during familiarity exploration sessions**. (A) Slide-window cross-Frequency phase-amplitude coupling concatenated in the four time-windows during Open-Field 2 exploration. Right panels show the modulation index during time for different pairs of coupling, comparison agains (0). (B) Shows the same as in (A), but for mPFC channels. (C) Shows the same as in (A), but for PAR channels. *p<0.05.

