## Supplementary Figures and Tables for "Beta2 oscillations in the hippocampal-cortical novelty detection circuit": Supplementary Figure 1.1.pdf

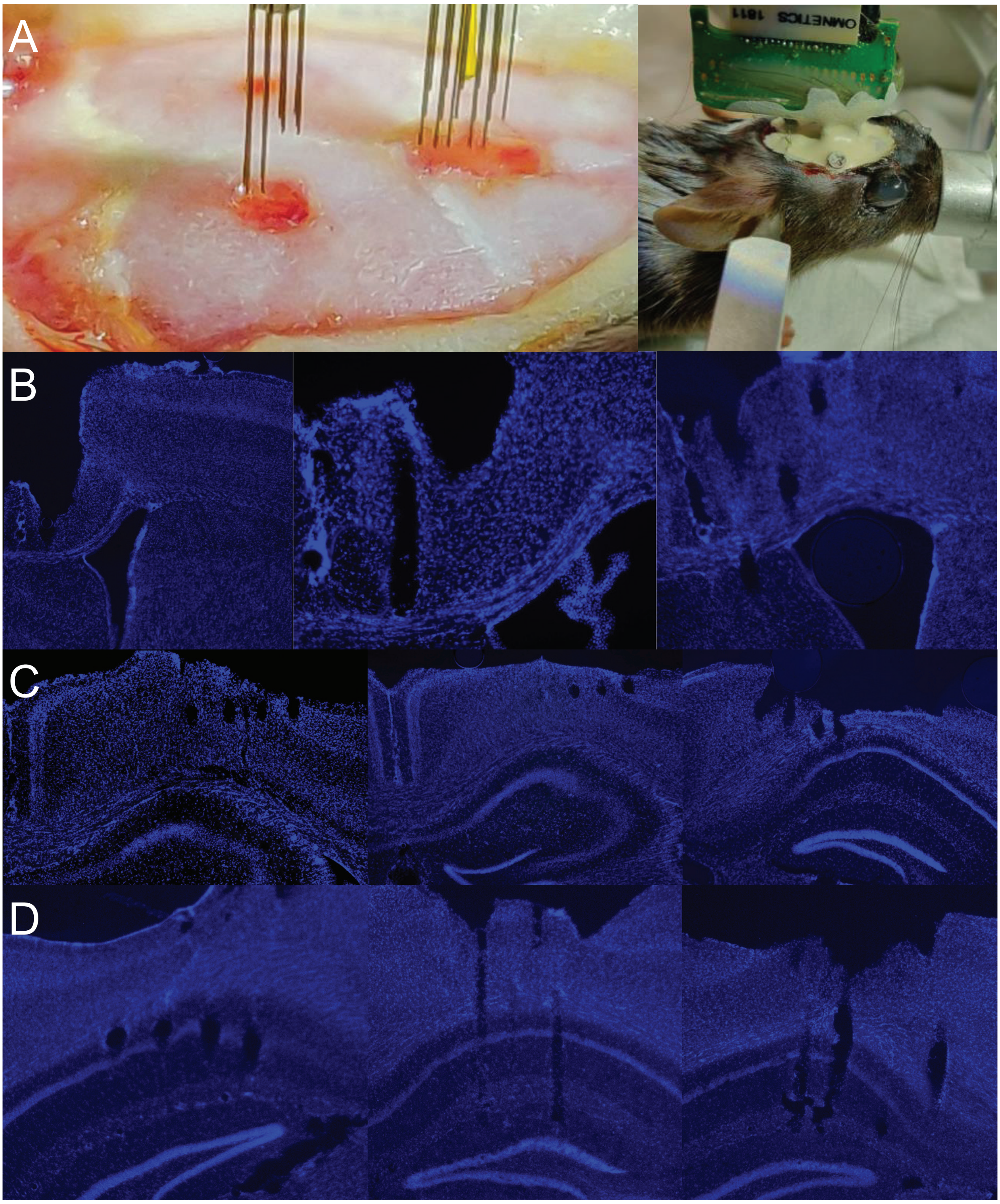


Supplementary Figure 1 – Electrodes positioning. (A) shows an example of electrode implant surgery. Left panel the two craniotomies for the electrodes of parietal cortex and hippocampus, and for the mid-frontal electrodes. Right panel shows the cap holding the implant to the head of the animal. (B) Examples of electrodes tracks found in mid-frontal cortex of the animals. (C) Example of electrodes tracks found in the parietal cortex. (D) Example of electrodes tracks found in the hippocampus.
