## Supplementary Figures and Tables for "Beta2 oscillations in the hippocampal-cortical novelty detection circuit": Supplementary Figure 2.pdf

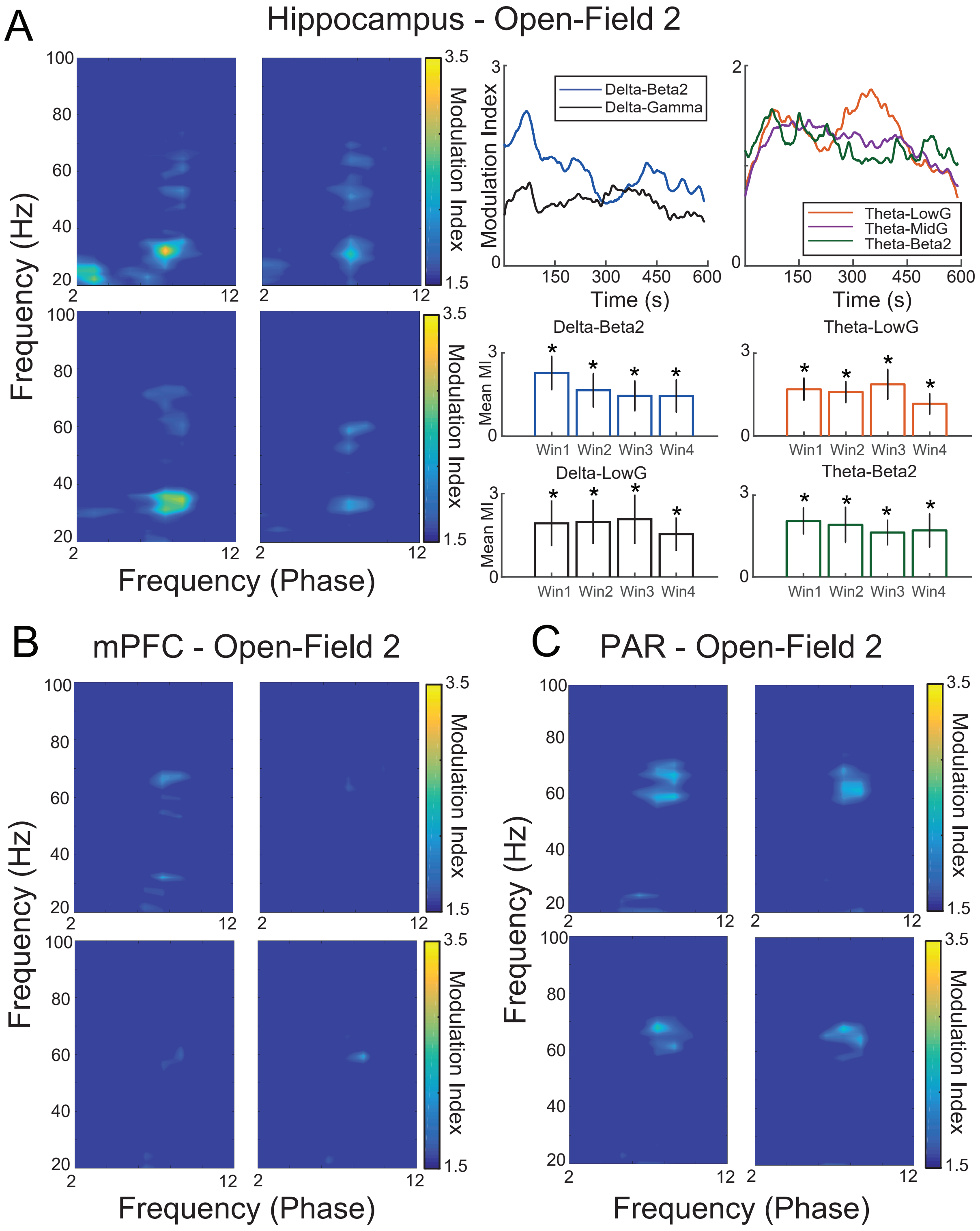


Supplementary Figure 2 – Cross-Frequency coupling during familiarity exploration sessions . (A) Slide-window cross-Frequency phase-amplitude coupling concatenated in the four time-windows during Open-Field 2 exploration. Right panels show the modulation index during time for different pairs of coupling, comparison agains (0). (B) Shows the same as in (A), but for mPFC channels. (C) Shows the same as in (A), but for PAR channels. *p<0.05.
