## Supplementary Figures and Tables for "Beta2 oscillations in the hippocampal-cortical novelty detection circuit": Table 1.pdf

| **Exploration session** | Metric | ANOVA Statistics |
|---|---|---|
| **All Sessions** | **Delta Power** time window comparison | $F_{8.5} = 1.01, p = 0.43$ |
| **All Sessions** | **Theta Power** time window comparison | $F_{8.5} = 1.76, p = 0.12$ |
| **All Sessions** | **Beta1 Power** time window comparison | $F_{8.5} = 0.69, p = 0.67$ |
| **All Sessions** | **Beta2 Power** time window comparison | $F_{8.5} = 2.75, p = 0.02$ |
| **All Sessions** | **LowG Power** time window comparison | $F_{8.5} = 2.18, p = 0.06$ |
| **Open Field 1** | **Beta2 Power** time window comparison | $F_{3.8} = 4.14, p = 0.01$ |
| **Open Field 2** | **Beta2 Power** time window comparison | $F_{3.8} = 6.21, p = 0.003$ |
| **Object 1** | **Beta2 Power** time window comparison | $F_{3.6} = 3.79, p = 0.02$; |
| **Object 2** | **Beta2 Power** time window comparison | $F_{3.6} = 3.03, p = 0.055$ |

Table 1 – Table of statistics related to figure 1. Descriptive statistics and comparisons with different sessions spectral power values and beta2 power time windows. Black: Significant p values, Red: non-significant p values.
