## Supplementary Figures and Tables for "Beta2 oscillations in the hippocampal-cortical novelty detection circuit": Table 2.pdf

| Exploration session (Brain Region) | Metric | Time-Window ANOVA Statistics | Paired t-test against 0 |
|---|---|---|---|
| **Open Field 1 (HC)** | **Delta/beta2** MI time window comparison | $F_{3,8}$ = 5.67, p = 0.004 | Win1 - $t_{2,8}$=3.13, p= 0.018 |
| | | | Win2 - $t_{2,8}$=3.56, p= 0.014 |
| | | | Win3 - $t_{2,8}$=3.82, p= 0.014 |
| | | | Win4 - $t_{2,8}$=2.91, p= 0.019 |
| **Open Field 1 (HC)** | **Delta/lowG** MI time window comparison | $F_{3,8}$ = 3.69, p = 0.025 | Win1 - $t_{2,8}$=3.72, p= 0.011 |
| | | | Win2 - $t_{2,8}$=4.03, p= 0.011 |
| | | | Win3 - $t_{2,8}$=3.08, p= 0.019 |
| | | | Win4 - $t_{2,8}$=2.87, p= 0.020 |
| **Open Field 1 (HC)** | **Theta/beta2** MI time window comparison | $F_{3,8}$ = 7.33, p = 0.001 | Win1 - $t_{2,8}$=4.97, p= 0.004 |
| | | | Win2 - $t_{2,8}$=4.17, p= 0.006 |
| | | | Win3 - $t_{2,8}$=3.47, p= 0.011 |
| | | | Win4 - $t_{2,8}$=1.74, p= 0.12 |
| **Open Field 1 (HC)** | **Theta/lowG** MI time window comparison | $F_{3,8}$ = 8.51, p = 0.0004 | Win1 - $t_{2,8}$=4.85, p= 0.002 |
| | | | Win2 - $t_{2,8}$=4.47, p= 0.002 |
| | | | Win3 - $t_{2,8}$=4.74, p= 0.002 |
| | | | Win4 - $t_{2,8}$=3.80, p= 0.005 |
| **Open Field 1 (HC)** | **Theta/midG** MI time window comparison | $F_{3,6}$ = 1.74, p = 0.18 | Win1 - $t_{2,8}$=3.01, p= 0.020 |
| | | | Win2 - $t_{2,8}$=2.96, p= 0.021 |
| | | | Win3 - $t_{2,8}$=4.74, p= 0.021 |
| | | | Win4 - $t_{2,8}$=2.93, p= 0.021 |
| **Object 1 (HC)** | **Delta/beta2** MI time window comparison | $F_{3,6}$= 2.53, p = 0.08 | Win1 - $t_{2,8}$=2.78, p= 0.027 |
| | | | Win2 - $t_{2,8}$=2.62, p= 0.034 |
| | | | Win3 - $t_{2,8}$=2.97, p= 0.027 |
| | | | Win4 - $t_{2,8}$=3.28, p= 0.013 |
| **Object 1 (HC)** | **Delta/lowG** MI time window comparison | $F_{3,6}$ = 1.44, p = 0.25 | Win1 - $t_{2,8}$=6.98, p= 0.001 |
| | | | Win2 - $t_{2,8}$=5.20, p= 0.005 |
| | | | Win3 - $t_{2,8}$=3.31, p= 0.021 |
| | | | Win4 - $t_{2,8}$=3.11, p= 0.017 |
| **Object 1 (HC)** | **Theta/beta2** MI time window comparison | $F_{3,6}$= 3.83, p = 0.02 | Win1 - $t_{2,8}$=3.75, p= 0.007 |
| | | | Win2 - $t_{2,8}$=3.07, p= 0.017 |
| | | | Win3 - $t_{2,8}$=2.28, p= 0.056 |
| | | | Win4 - $t_{2,8}$=3.13, p= 0.016 |
| **Object 1 (HC)** | **Theta/lowG** MI time window comparison | $F_{3,6}$= 4.13, p = 0.01; | Win1 - $t_{2,8}$=5.36, p= 0.001 |
| | | | Win2 - $t_{2,8}$=4.24, p= 0.003 |
| | | | Win3 - $t_{2,8}$=5.11, p= 0.001 |
| | | | Win4 - $t_{2,8}$=3.61, p= 0.008 |
| **Object 1 (HC)** | **Theta/midG** MI time window comparison | F3,6 = 0.40, p = 0.74 | Win1 - $t_{2,8}$=4.03, p= 0.002 |
| | | | Win2 - $t_{2,8}$=2.37, p= 0.030 |
| | | | Win3 - $t_{2,8}$=2.51, p= 0.028 |
| | | | Win4 - $t_{2,8}$=3.35, p= 0.008 |

Table 2 – Table of statistics related to figure 2. Descriptive statistics and comparisons with different MI time window analysis and the comparison with null hypothesis 0. Black: Significant p values, Red: non-significant p values.
