## Supplementary Figures and Tables for "Beta2 oscillations in the hippocampal-cortical novelty detection circuit": Table 3.pdf

| | *OpenField 1 Session* | | |
|---|---|---|---|
| *Exploration session* | Metric | Time-Window ANOVA Statistics | Paired t-test against 0 |
| ***Open Field 1 mPFC*** | **Beta2 Power** time-window comparison | ($F_{3,8}$ = 6.53, p = 0.002) | |
| ***Open Field 1 mPFC*** | **Delta/beta2** MI time window comparison | $F_{3,8}$ = 11.4, p = 0.00007 | Win1 - $t_{2,8}$=4.09, p= 0.003 |
| | | | Win2 - $t_{2,8}$=3.56, p= 0.026 |
| | | | Win3 - $t_{2,8}$=3.82, p= 0.237 |
| | | | Win4 - $t_{2,8}$=2.91, p= 0.501 |
| ***Open Field 1 mPFC*** | **Delta/lowG** MI time window comparison | $F_{3,8}$ = 4.85, p = 0.0088 | Win1 - $t_{2,8}$=2.71, p= 0.026 |
| | | | Win2 - $t_{2,8}$=4.03, p= 0.033 |
| | | | Win3 - $t_{2,8}$=3.08, p= 0.22 |
| | | | Win4 - $t_{2,8}$=2.87, p= 0.67 |
| ***Open Field 1 mPFC*** | **Theta/beta2** MI time window comparison | $F_{3,8}$ = 11.42, p > 0.0001 | Win1 - $t_{2,8}$=3.81, p= 0.005 |
| | | | Win2 - $t_{2,8}$=3.01, p= 0.016 |
| | | | Win3 - $t_{2,8}$=1.03, p= 0.32 |
| | | | Win4 - $t_{2,8}$=0.13, p= 0.89 |
| ***Open Field 1 mPFC*** | **Theta/lowG** MI time window comparison | $F_{3,8}$ = 5.06, p = 0.007 | Win1 - $t_{2,8}$=2.97, p= 0.017 |
| | | | Win2 - $t_{2,8}$=2.98, p= 0.017 |
| | | | Win3 - $t_{2,8}$=2.52, p= 0.035 |
| | | | Win4 - $t_{2,8}$=1.75, p= 0.116 |
| ***Open Field 1 mPFC*** | **Theta/midG** MI time window comparison | $F_{3,8}$ = 2.72, p = 0.066 | Win1 - $t_{2,8}$=2.36, p= 0.045 |
| | | | Win2 - $t_{2,8}$=2.48, p= 0.038 |
| | | | Win3 - $t_{2,8}$=2.63, p= 0.030 |
| | | | Win4 - $t_{2,8}$=2.88, p= 0.020 |
| ***Open Field 1 PAR*** | **Beta2 Power** time-window comparison | ($F_{3,8}$ = 1.07, p = 0.38) | |
| ***Open Field 1 PAR*** | **Delta/beta2** MI time window comparison | $F_{3,6}$= 14.62, p > 0.0001 | Win1 - $t_{2,8}$=3.35, p= 0.006 |
| | | | Win2 - $t_{2,8}$=2.65, p= 0.058 |
| | | | Win3 - $t_{2,8}$=2.20, p= 0.213 |
| | | | Win4 - $t_{2,8}$=1.36, p= 0.746 |
| ***Open Field 1 PAR*** | **Delta/lowG** MI time window comparison | $F_{3,6}$ = 8.09, p = 0.0006 | Win1 - $t_{2,8}$=2.67, p= 0.028 |
| | | | Win2 - t2,8=1.80, p=0.108 |
| | | | Win3 - $t_{2,8}$=3.31, p= 0.122 |
| | | | Win4 - $t_{2,8}$=0.09, p= 0.929 |
| ***Open Field 1 PAR*** | **Theta/beta2** MI time window comparison | $F_{3,6}$= 21.03, p > 0.0001 | Win1 - $t_{2,8}$=5.15,p=0.0009 |
| | | | Win2 - $t_{2,8}$=3.33, p= 0.010 |
| | | | Win3 - $t_{2,8}$=1.77, p= 0.113 |
| | | | Win4 - $t_{2,8}$=-0.20, p=0.845 |
| ***Open Field 1*** | | $F_{3,6}$= 8.56, p = 0.0004; | Win1 - $t_{2,8}$=3.12, p= 0.014 |

| | | | |
|---|---|---|---|
| ***PAR*** | **Theta/lowG** MI time window comparison | | Win2 - $t_{2,8}$=2.20, p= 0.058<br>Win3 - $t_{2,8}$=1.80, p= 0.109<br>Win4 - $t_{2,8}$=1.43, p= 0.189 |
| ***Open Field 1 PAR*** | **Theta/midG** MI time window comparison | F3,6 = 3.98, p = 0.019 | Win1 - $t_{2,8}$=3.26, p= 0.011<br>Win2 - $t_{2,8}$=3.39, p= 0.009<br>Win3 - $t_{2,8}$=3.49, p= 0.008<br>Win4 - $t_{2,8}$=3.28, p= 0.011 |
| | ***Object 1 Session*** | | |
| ***Exploration session*** | **Metric** | **Time-Window ANOVA Statistics** | **Paired t-test against 0** |
| ***Object 1 mPFC*** | **Beta2 Power** time-window comparison | ($F_{3,6}$ = 2.4, p = 0.10) | |
| ***Object 1 mPFC*** | **Delta/beta2** MI time window comparison | $F_{3,6}$ = 3.97, p = 0.021 | Win1 - $t_{2,7}$=2.65, p= 0.040<br>Win2 - $t_{2,7}$=2.14, p= 0.094<br>Win3 - $t_{2,7}$=0.82, p= 0.253<br>Win4 - $t_{2,7}$=2.13, p= 0.034 |
| ***Object 1 mPFC*** | **Delta/lowG** MI time window comparison | $F_{3,6}$ = 0.99, p = 0.41 | Win1 - $t_{2,7}$=2.18, p= 0.065<br>Win2 - $t_{2,7}$=2.28, p= 0.056<br>Win3 - $t_{2,7}$=2.14, p= 0.068<br>Win4 - $t_{2,7}$=2.73, p= 0.029 |
| ***Object 1 mPFC*** | **Theta/beta2** MI time window comparison | $F_{3,6}$ = 5.45, p > 0.006 | Win1 - $t_{2,7}$=4.41, p= 0.003<br>Win2 - $t_{2,7}$=2.70, p= 0.027<br>Win3 - $t_{2,7}$=1.68, p= 0.144<br>Win4 - $t_{2,7}$=2.13, p= 0.070 |
| ***Object 1 mPFC*** | **Theta/lowG** MI time window comparison | $F_{3,6}$ = 1.21, p = 0.330 | Win1 - $t_{2,8}$=2.97, p= 0.040<br>Win2 - $t_{2,8}$=2.98, p= 0.046<br>Win3 - $t_{2,8}$=2.52, p= 0.033<br>Win4 - $t_{2,8}$=1.75, p= 0.032 |
| ***Object 1 mPFC*** | **Theta/midG** MI time window comparison | $F_{3,6}$ = 1.45, p = 0.254 | Win1 - $t_{2,8}$=2.58, p= 0.037<br>Win2 - $t_{2,8}$=1.86, p= 0.104<br>Win3 - $t_{2,8}$=2.05, p= 0.079<br>Win4 - $t_{2,8}$=2.88, p= 0.021 |
| ***Object 1 PAR*** | **Beta2 Power** time-window comparison | ($F_{3,6}$ = 1.16, p = 0.35) | |
| ***Object 1 PAR*** | **Delta/beta2** MI time window comparison | $F_{3,6}$= 2.95, p = 0.060 | Win1 - $t_{2,8}$=3.55, p= 0.012<br>Win2 - $t_{2,8}$=2.65, p= 0.255<br>Win3 - $t_{2,8}$=2.20, p= 0.323<br>Win4 - $t_{2,8}$=1.36, p= 0.941 |
| ***Object 1 PAR*** | **Delta/lowG** MI time window comparison | $F_{3,6}$ = 1.79, p = 0.18 | Win1 - $t_{2,8}$=2.16, p= 0.073<br>Win2 - $t_{2,8}$=1.80, p= 0.308<br>Win3 - $t_{2,8}$=3.31, p= 0.106<br>Win4 - $t_{2,8}$=0.09, p= 0.880 |

| | | | |
|---|---|---|---|
| ***Object 1 PAR*** | **Theta/beta2** MI time window comparison | $F_{3,6}$= 0.57, p = 0.64 | Win1 - $t_{2,8}$=2.26, p= 0.063 |
| | | | Win2 - $t_{2,8}$=1.15, p= 0.290 |
| | | | Win3 - $t_{2,8}$=2.80, p= 0.030 |
| | | | Win4 - $t_{2,8}$=1.19, p= 0.278 |
| ***Object 1 PAR*** | **Theta/lowG** MI time window comparison | $F_{3,6}$= 0.58, p = 0.63 | Win1 - $t_{2,8}$=2.26, p= 0.173 |
| | | | Win2 - $t_{2,8}$=1.15, p= 0.451 |
| | | | Win3 - $t_{2,8}$=2.80, p= 0.079 |
| | | | Win4 - $t_{2,8}$=1.19, p= 0.087 |
| ***Object 1 PAR*** | **Theta/midG** MI time window comparison | $F_{3,6}$ = 1.37, p = 0.28 | Win1 - $t_{2,8}$=3.26, p= 0.127 |
| | | | Win2 - $t_{2,8}$=3.39, p= 0.221 |
| | | | Win3 - $t_{2,8}$=3.49, p= 0.075 |
| | | | Win4 - $t_{2,8}$=3.28, p= 0.100 |

Table 3 – Table of statistics related to figure 6. Descriptive statistics and comparisons with different MI time window analysis and the comparison with null hypothesis 0. Black: Significant p values, Red: non-significant p values.
