## Supplementary Figures and Tables for "Beta2 oscillations in the hippocampal-cortical novelty detection circuit": Table 4.pdf

| Exploration session | Metric | Time-Window ANOVA Statistics | Paired t-test against 0 |
|---|---|---|---|
| **Open Field 2 (HC)** | **Delta/beta2** MI time window comparison | $F_{3,8}$ = 1.12, p = 0.359 | Win1 - $t_{2,8}$=3.25, p= 0.011 |
| | | | Win2 - $t_{2,8}$=2.74, p= 0.025 |
| | | | Win3 - $t_{2,8}$=2.76, p= 0.024 |
| | | | Win4 - $t_{2,8}$=2.82, p= 0.022 |
| **Open Field 2 (HC)** | **Delta/lowG** MI time window comparison | $F_{3,8}$ = 0.22, p = 0.223 | Win1 - $t_{2,8}$=2.85, p= 0.021 |
| | | | Win2 - $t_{2,8}$=2.76, p= 0.024 |
| | | | Win3 - $t_{2,8}$=2.59, p= 0.031 |
| | | | Win4 - $t_{2,8}$=3.81, p= 0.005 |
| **Open Field 2 (HC)** | **Theta/beta2** MI time window comparison | $F_{3,8}$ = 0.41, p = 0.74 | Win1 - $t_{2,8}$=6.68, p> 0.001 |
| | | | Win2 - $t_{2,8}$=3.45, p= 0.008 |
| | | | Win3 - $t_{2,8}$=4.16, p= 0.003 |
| | | | Win4 - $t_{2,8}$=3.04, p= 0.015 |
| **Open Field 2 (HC)** | **Theta/lowG** MI time window comparison | $F_{3,8}$ = 2.44, p = 0.088 | Win1 - $t_{2,8}$=4.77, p= 0.001 |
| | | | Win2 - $t_{2,8}$=4.35, p= 0.002 |
| | | | Win3 - $t_{2,8}$=4.32, p= 0.002 |
| | | | Win4 - $t_{2,8}$=4.08, p= 0.003 |
| **Open Field 2 (HC)** | **Theta/midG** MI time window comparison | $F_{3,8}$ = 1.44, p = 0.25 | Win1 - $t_{2,8}$=2.93, p= 0.019 |
| | | | Win2 - $t_{2,8}$=2.92, p= 0.019 |
| | | | Win3 - $t_{2,8}$=2.96, p= 0.018 |
| | | | Win4 - $t_{2,8}$=2.79, p= 0.023 |
| **Open Field 2 (mPFC)** | **Delta/beta2** MI time window comparison | $F_{3,8}$= 0.86, p = 0.475 | Win1 - $t_{2,8}$=1.83, p= 0.109 |
| | | | Win2 - $t_{2,8}$=1.38, p= 0.208 |
| | | | Win3 - $t_{2,8}$=2.40, p= 0.046 |
| | | | Win4 - $t_{2,8}$=1.71, p= 0.129 |
| **Open Field 2 (mPFC)** | **Delta/lowG** MI time window comparison | $F_{3,8}$ = 0.88, p = 0.464 | Win1 - $t_{2,8}$=3.22, p= 0.042 |
| | | | Win2 - $t_{2,8}$=1.88, p= 0.141 |
| | | | Win3 - $t_{2,8}$=2.02, p= 0.174 |
| | | | Win4 - $t_{2,8}$=1.44, p= 0.220 |
| **Open Field 2 (mPFC)** | **Theta/beta2** MI time window comparison | $F_{3,8}$= 0.28, p = 0.83 | Win1 - $t_{2,8}$=2.65, p= 0.032 |
| | | | Win2 - $t_{2,8}$=2.52, p= 0.039 |
| | | | Win3 - $t_{2,8}$=3.23, p= 0.014 |
| | | | Win4 - $t_{2,8}$=2.50, p= 0.040 |
| **Open Field 2 (mPFC)** | **Theta/lowG** MI time window comparison | $F_{3,8}$= 0.77, p = 0.510; | Win1 - $t_{2,8}$=3.78, p= 0.006 |
| | | | Win2 - $t_{2,8}$=3.02, p= 0.019 |
| | | | Win3 - $t_{2,8}$=2.21, p= 0.062 |
| | | | Win4 - $t_{2,8}$=2.59, p= 0.035 |
| **Open Field 2 (mPFC)** | **Theta/midG** MI time window comparison | $F_{3,8}$=0.36, p = 0.781 | Win1 - $t_{2,8}$=4.03, p= 0.133 |
| | | | Win2 - $t_{2,8}$=2.37, p= 0.061 |
| | | | Win3 - $t_{2,8}$=2.51, p= 0.035 |
| | | | Win4 - $t_{2,8}$=3.35, p= 0.063 |
| **Open Field 2 (PAR)** | **Delta/beta2** MI time window comparison | $F_{3,8}$= 0.50, p = 0.683 | Win1 - $t_{2,8}$=1.47, p= 0.191 |
| | | | Win2 - $t_{2,8}$=1.57, p= 0.166 |

| | | | |
|---|---|---|---|
| | | | Win3 - $t_{2,8}$=2.77, p= 0.032 |
| | | | Win4 - $t_{2,8}$=2.46, p= 0.048 |
| **Open Field 2 (PAR)** | **Delta/lowG** MI time window comparison | $F_{3,8}$ = 0.63, p = 0.599 | Win1 - $t_{2,8}$=1.84, p= 0.114 |
| | | | Win2 - $t_{2,8}$=0.48, p= 0.642 |
| | | | Win3 - $t_{2,8}$=025, p= 0.810 |
| | | | Win4 - $t_{2,8}$=1.39, p= 0.213 |
| **Open Field 2 (PAR)** | **Theta/beta2** MI time window comparison | $F_{3,8}$= 0.39, p = 0.755 | Win1 - $t_{2,8}$=2.62, p= 0.039 |
| | | | Win2 - $t_{2,8}$=2.31, p= 0.059 |
| | | | Win3 - $t_{2,8}$=2.98, p= 0.024 |
| | | | Win4 - $t_{2,8}$=2.41, p= 0.052 |
| **Open Field 2 (PAR)** | **Theta/lowG** MI time window comparison | $F_{3,8}$= 1.03, p = 0.40; | Win1 - $t_{2,8}$=1.44, p= 0.198 |
| | | | Win2 - $t_{2,8}$=2.11, p= 0.078 |
| | | | Win3 - $t_{2,8}$=1.60, p= 0.159 |
| | | | Win4 - $t_{2,8}$=1.95, p= 0.098 |
| **Open Field 2 (PAR)** | **Theta/midG** MI time window comparison | $F_{3,8}$=0.21, p = 0.881 | Win1 - $t_{2,8}$=1.74, p= 0.132 |
| | | | Win2 - $t_{2,8}$=2.10, p= 0.079 |
| | | | Win3 - $t_{2,8}$=2.07, p= 0.083 |
| | | | Win4 - $t_{2,8}$=2.14, p= 0.075 |

Table 4 – Table of statistics related to the supplementary figure 2. Descriptive statistics and comparisons with different MI time window analysis and the comparison with null hypothesis 0. Black: Significant p values, Red: non-significant p values.
